# Microglia integration into midbrain organoids – origin matters

**DOI:** 10.64898/2026.09.22.753683

**Authors:** Marloes Verkerke, Angélica Maria Sabogal-Guáqueta, Saloua Zekri, Anne-Marie van Dam, Helga E. de Vries, Vanessa Donega

**Author notes:** Corresponding author: Vanessa Donega –. Shared first author (Marloes Verkerke, Angélica Maria Sabogal-Guáqueta). Shared last author (Helga E. de Vries, Vanessa Donega).

## Abstract

Midbrain organoids (MOs) can be powerful tools to study brain diseases, yet one shortcoming is the lack of microglia due to their mesodermal developmental origin. Previous studies have incorporated iPSC-derived microglia (iMG), but their immature phenotype restricts the applicability to model age-related neuroinflammatory processes relevant to diseases such as Parkinson’s Disease (PD). To address this, we have incorporated MOs with human postmortem PD microglia, iPSC-derived microglial progenitors (iMP), and iMG. Postmortem microglia had a higher infiltration capacity resulting in more microglia per organoid, while iMP had a higher migration capacity reaching the core of the organoid. Compared to iPSC-derived cells, more infiltrated postmortem PD microglia were HLA Class II positive and CD68 positive. This novel approach of integrating postmortem microglia into organoids opens new avenues for modelling age-related neuroinflammatory diseases. Ongoing methodological advancements like our study will offer the field a valuable tool for studying human microglia biology in both health and disease.

**Highlights:**

- Postmortem microglia have a higher infiltration capacity resulting in more microglia per organoid
- iPSC-derived microglial progenitors could have a higher migration capacity to reach the organoid core
- More postmortem microglia express HLA Class II and CD68 than iPSC-derived cells
- P2RY12 and IBA1 expression is higher in iPSC-derived microglial progenitors

**eTOC:** Verkerke and colleagues explore a novel approach of integrating postmortem microglia from Parkinson’s Disease donors into midbrain organoids. Comparison to integration of iPSC-derived microglia and microglial progenitors reveals a higher infiltration capacity of postmortem microglia with an elevated presence of inflammatory markers HLA Class II and CD68. This innovative approach paves the way for new insights into age-related neuroinflammatory processes using organoid models.

## INTRODUCTION

Parkinson’s disease (PD) is the second most common neurodegenerative disorder, characterized by the progressive loss of dopaminergic neurons in the substantia nigra and the pathological accumulation of α-synuclein, particularly within Lewy Bodies (Elbaz et al., 2016). This progressive neuronal loss results in motor dysfunction and cognitive decline, yet current treatments remain purely symptomatic as no clinically approved disease-modifying therapies exist (Aarsland et al., 2021; Panicker et al., 2021). Neuroinflammation is increasingly recognized as an important component of PD and is mediated by microglia, the brain’s resident immune cells (Araújo et al., 2022). Microglia are currently understood to exist along a spectrum of functional states, dynamically adapting to changes in their environment (Gao et al., 2023; Vidal-Itriago et al., 2022). In their homeostatic state, microglia exhibit a highly ramified morphology that allows them to continuously monitor and interact with the surrounding microenvironment. In response to inflammatory or pathological stimuli, microglia can transition into an active, pro-inflammatory profile. This profile is characterized by process retraction and the release of pro-inflammatory cytokines and chemokines that amplify the immune response by facilitating the recruitment of peripheral immune cells (Gao *et al*., 2023). In PD, however, this microglial activation becomes sustained, contributing to chronic neuroinflammation (Isik et al., 2023). Postmortem studies of human PD brains have demonstrated the presence of reactive microglia in affected regions, such as the substantia nigra and striatum, where microglia display an activated morphology and express pro-inflammatory markers like HLA-DR (Imamura et al., 2003; McGeer et al., 1988). Moreover, already in early PD, increased expression of Toll-like receptor 2 was observed in the hippocampus and substantia nigra (Doorn et al., 2014; Kim et al., 2013). Complementing these observations, pharmacological depletion of mouse microglia was shown to lead to reduced α-synuclein accumulation and dopaminergic neurodegeneration (Thi Lai et al., 2024). However, mechanistic studies of microglia-mediated neuroinflammation in human-derived models are still limited, underscoring the need for more physiologically relevant in vitro systems for investigating microglial contributions to PD pathology.

The human brain’s complex structure and vast cellular network present a major challenge for disease modelling. Advancements in stem cell technologies have enabled the development of 3D brain organoids from human induced pluripotent stem cells (iPSCs), including region-specific models that mimic distinct areas of the human brain (Paşca, 2018). Conventional brain organoid protocols generate tissue of ectodermal origin and therefore lack microglia. Besides mediating neuroinflammatory responses, microglia are essential for shaping neural circuits through synaptic pruning and clearing apoptotic cells in the brain (Isik *et al*., 2023; Paolicelli et al., 2011; Sierra et al., 2013). Their absence from brain organoid models thus limits both the cellular and functional complexity of these systems, and prevents the study of neuroinflammation in disorders such as PD. To overcome this absence, previous studies have integrated iPSC-derived microglia into brain organoids or manipulated culture conditions to enable the innate development of microglia within brain organoids. Including microglia within brain organoids allows interaction with neurons and other glial cells in a physiologically relevant environment, capturing complex behaviors like migration, synaptic pruning, and immune responses that are absent in 2D cultures (Abud et al., 2017; Fagerlund et al., 2021; Ormel et al., 2018; Wenzel et al., 2024).

Midbrain organoids (MOs) have emerged as a valuable model for studying PD, as they generate dopaminergic neurons in a spatially organized manner, along astrocytes and oligodendrocytes (Monzel et al., 2017; Smits et al., 2019). Most importantly, MOs with PD-related genetic alterations have been shown to model α-synuclein aggregation as well as the loss of dopaminergic neurons (Kim et al., 2019; Smits *et al*., 2019). Yet, like other ectoderm-derived brain organoids, MOs lack microglia. Sabate et al., 2022 overcame this limitation by integrating iPSC-derived microglia and demonstrated that these microglia influence neuronal development and increase the applicability of organoid models for studying neurodegenerative diseases (Sabate-Soler et al., 2022).

In the present study, we show for the first time that microglia derived from postmortem PD patients integrate into MOs within just a few days. Within the organoid environment, we compared the infiltration capacity and phenotypes of the postmortem PD microglia with iPSC-derived microglial progenitors (iMP) and iPSC-derived microglia (iMG). Postmortem PD microglia exhibited an approximately three-fold higher infiltration capacity compared to both iPSC-derived cells. On the other hand, more iPSC-derived cells compared to postmortem microglia reached the organoid core indicating a higher migration capacity. Phenotypic characterization of infiltrated microglia revealed a high proportion of postmortem PD microglia expressing reactive markers HLA Class II and CD68, whereas a higher proportion of iMP expressed homeostatic markers P2RY12 and IBA1.

Taken together, our study is the first to our knowledge to report the infiltration capacity of postmortem PD microglia into MOs and to describe the phenotypic differences between infiltrated microglia from postmortem or iPSC origin. Similar to our study, ongoing methodological advancements in incorporating microglia into organoids will offer the field a valuable tool for studying human microglia biology in both health and disease.

## RESULTS

### Postmortem microglia show greater infiltration than iPSC-derived microglial progenitors and microglia

To determine whether microglia from different origins spontaneously infiltrate organoids, 5-week-old MOs were cultured with a suspension of fluorescently-labelled (CFSE) microglia for two weeks (**Fig. 1A**). We compared the infiltration capacities across three groups: human postmortem PD microglia (PD), iPSC-derived microglial progenitors (iMP), and iPSC-derived microglia (iMG). Growth factors were added to the iMP group to ensure maturation of microglia.

**Figure 1.**
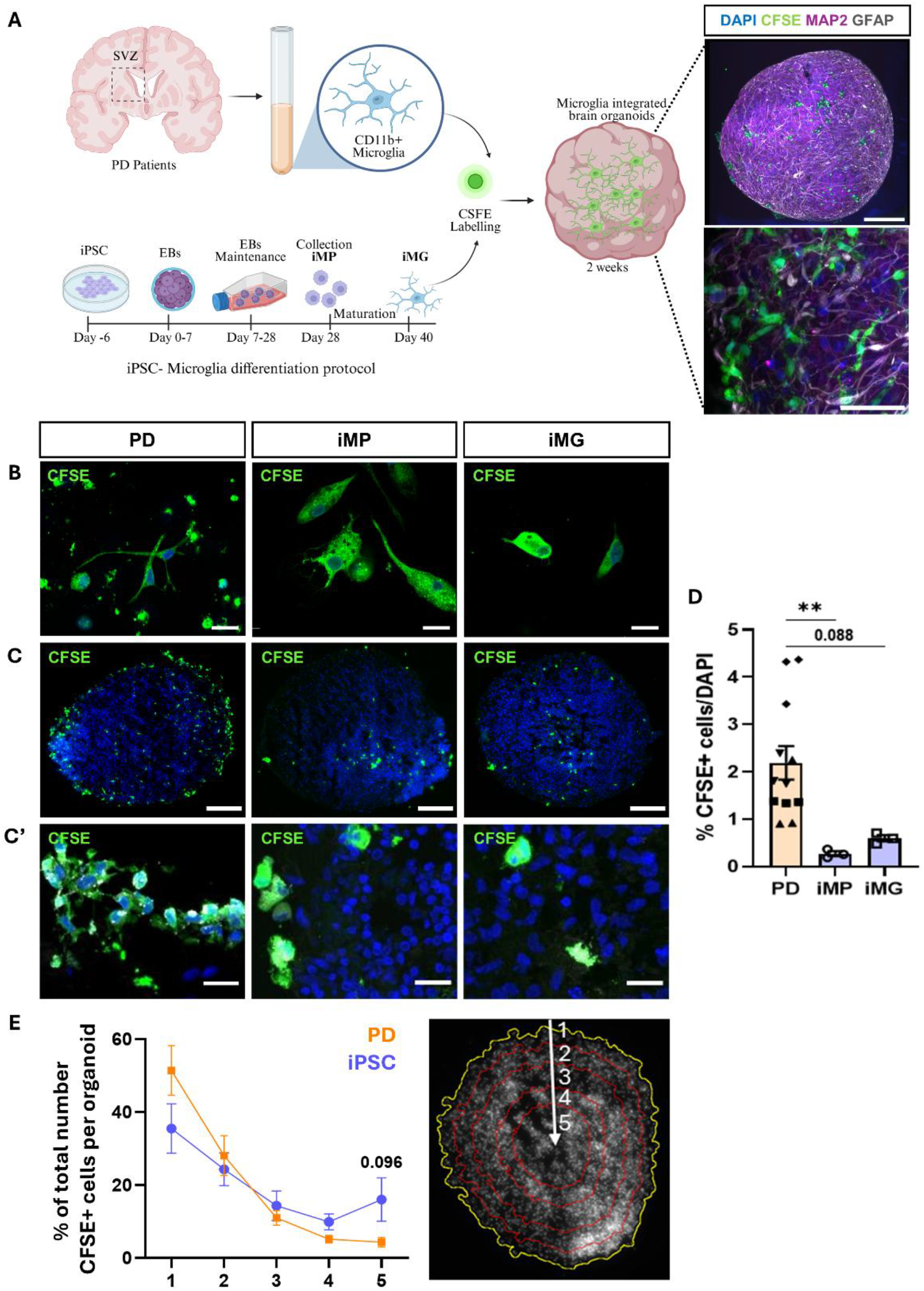
Incorporation of postmortem and iPSC-derived microglia into midbrain organoids. **A** Experimental overview and representative image of whole midbrain organoid (MO; DIV50) consisting of MAP2+ neurons (magenta) and GFAP+ astrocytes (grey) with infiltrated CFSE+ microglia (green). Nuclei (DAPI) in blue. Scale bar top = 200 µm. Scale bar bottom = 50 µm. **B** Representative images of monoculture (24 hours) of CFSE+ microglia from postmortem PD tissue (PD), iPSC-derived microglia progenitor (iMP), and iPSC-derived microglia (iMG). Scale bars = 20 µm. **C** Representative images of MOs with infiltrated CFSE+ microglia. Scale bars = 200 µm. **C’** High magnification of infiltrated CFSE+ microglia. Scale bar = 20 µm. **D** Quantification of infiltrated CFSE+ microglia as percentage of total nuclei per organoid section. Each symbol represents an organoid. Kruskal-Wallis with Dunn’s multiple comparisons test was performed. ^**^ p_adj_ ≤ 0.01. **E** Quantification of infiltrated CFSE+ microglia from the edge of the organoid to the core of the organoid as percentage of total number of infiltrated CFSE+ microglia. Multiple unpaired t-tests with Holm-Šidák method were performed. All data shown as mean ±SEM.

Successful labelling with CFSE of microglia from all three origins was confirmed in monoculture (**Fig 1B**). Microglia from all three origins – PD, iMP, and iMG – spontaneously infiltrated the MOs (**Fig. 1C-C’**). Significantly more PD microglia (2.18% ± 0.35%) infiltrated the MOs compared to the iMP (0.27% ± 0.05%; p_adj_ = 0.007) and iMG (0.59% ± 0.07%; p_adj_ = 0.088) (**Fig. 1D**). Within the iPSC group, there was no difference between iMP and iMG. Finally, we quantified how far into the MO the microglia infiltrated as a measure of migration capacity. For this analysis, we divided each organoid section into shells with the outer rim as shell 1 and the core as shell 5 and subsequently compared postmortem and iPSC microglia (iMP and iMG combined) (**Fig. 1E**). Of the total number of infiltrated microglia, the majority of microglia resided in the outer rims of the organoid which was similar in both groups (PD_shell1_ 51.39% ± 6.86%; PD_shell2_ 28.08% ± 5.49%; iPSC_shell1_ 35.49% ± 6.75%; iPSC_shell2_ 24.33% ± 4.49%). We did find a trend implicating more iPSC-derived microglia (16.0% ± 6.0%) compared to postmortem PD microglia (4.36% ± 1.33%) reached the organoid core (p_adj_ = 0.096). In conclusion, postmortem microglia exhibited higher infiltration capacity compared to iPSC microglia, whereas iPSC microglia could have higher migration capacity.

### Increased number of P2RY12 and IBA1 positive iPSC-derived microglia progenitors

Phenotypic characteristics of the microglia within the organoid environment were assessed using P2RY12 and IBA1 as homeostatic markers and HLA Class II and CD68 as reactive markers. The number of positive cells was corrected for the number of total CFSE+ cells as a correction for the demonstrated higher infiltration capacity of PD microglia compared to iMP and iMG.

P2RY12+ cells were observed in all three groups (**Fig. 2**). Of the infiltrated cells, significantly more iMP (3.4 ± 1.08) showed P2RY12 immunoreactivity compared to PD microglia (0.97 ± 0.12; p_adj_ = 0.0008) and iMG (1.37 ± 0.43; p_adj_ = 0.019). A similar pattern was observed for IBA1, showing IBA1+ cells in all three groups with significantly more IBA1+ iMP (3.64 ± 0.38) compared to PD (1.64 ± 0.17; p_adj_ = 0.0002) and iMG (1.60 ± 0.12; p_adj_ = 0.0013) (**Fig. 3**).

**Figure 2.**
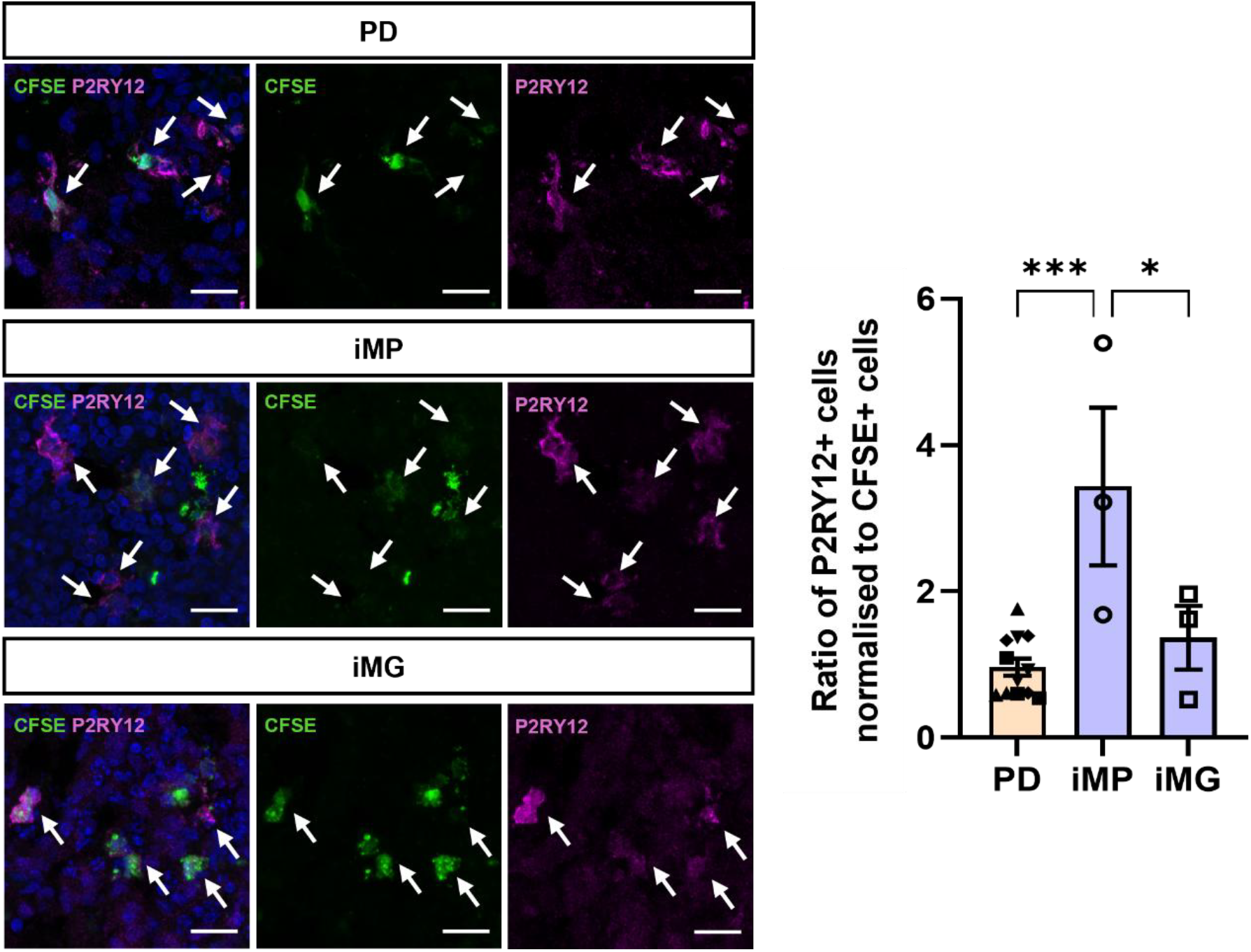
Expression of P2RY12 in microglia infiltrated into midbrain organoids. Representative images of infiltrated CFSE+ microglia from postmortem PD tissue (PD), iPSC-derived microglia progenitors (iMG), and iPSC-derived microglia (iMG) stained for P2RY12. White arrows show examples of P2RY12+ cells. Nuclei (DAPI) in blue. Scale bars = 50 µm. Quantification of the ratio of infiltrated CFSE+ microglia that express P2RY12. An ordinary one-way ANOVA with Tukey’s multiple comparisons test was performed. Each symbol represents an organoid. Data shown as mean ± SEM. ^*^ p_adj_ ≤ 0.05 ^**^ p_adj_ ≤ 0.01 ^***^ Padj ≤ 0.001.

**Figure 3.**
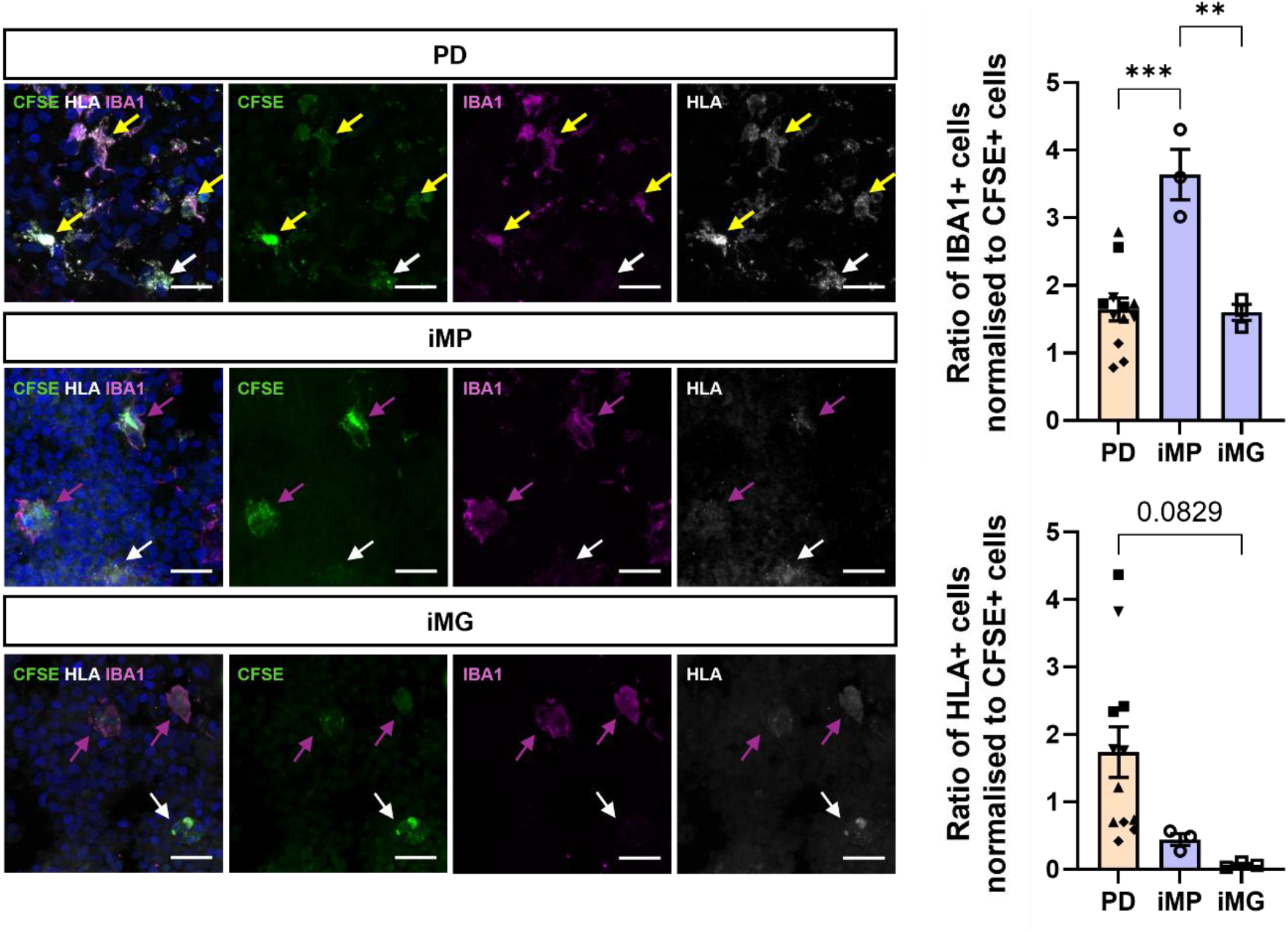
Expression of IBA1 and HLA in microglia infiltrated into midbrain organoids. Representative images of infiltrated CFSE+ microglia from postmortem PD tissue (PD), iPSC-derived microglia progenitors (iMG), and iPSC-derived microglia (iMG) stained for IBA1 and HLA. Yellow arrows show examples of IBA1+ and HLA+ double positive cells. White arrows show examples of HLA+ cells and magenta arrows show examples of IBA1+ cells. Nuclei (DAPI) in blue. Scale bars = 50 µm. Quantification of the ratio of infiltrated CFSE+ microglia that express IBA1 or HLA. An ordinary one-way ANOVA with Tukey’s multiple comparisons test was performed. Each symbol represents an organoid. Data shown as mean ±SEM. ^*^ p_adj_ ≤ 0.05 ^**^ p_adj_ ≤ 0.01 ^***^ p_adj_ ≤ 0.001.

A different pattern was observed for the reactive microglia markers. Although HLA+ cells were present in all three groups, a trend towards increased number of HLA+ cells in the PD group (1.74 ± 0.38) compared to iMG was observed (0.06 ± 0.02; p_adj_ = 0.0829) whereas no difference was found compared to iMP (0.44 ± 0.09) (**Fig 3**). Additionally, the HLA signal intensity appeared higher in PD with strong expression throughout the cell bodies and branches of the microglia. Lysosomal protein CD68 was mainly present in the cell bodies and a similar expression pattern was observed as for HLA. Significantly more cells showed CD68 immunoreactivity in the PD group (0.68 ± 0.12) compared to iMG (0.29 ± 0.07; p_adj_ = 0.041) with no difference to iMP (0.44 ± 0.002) (**Fig. 4**).

**Figure 4.**
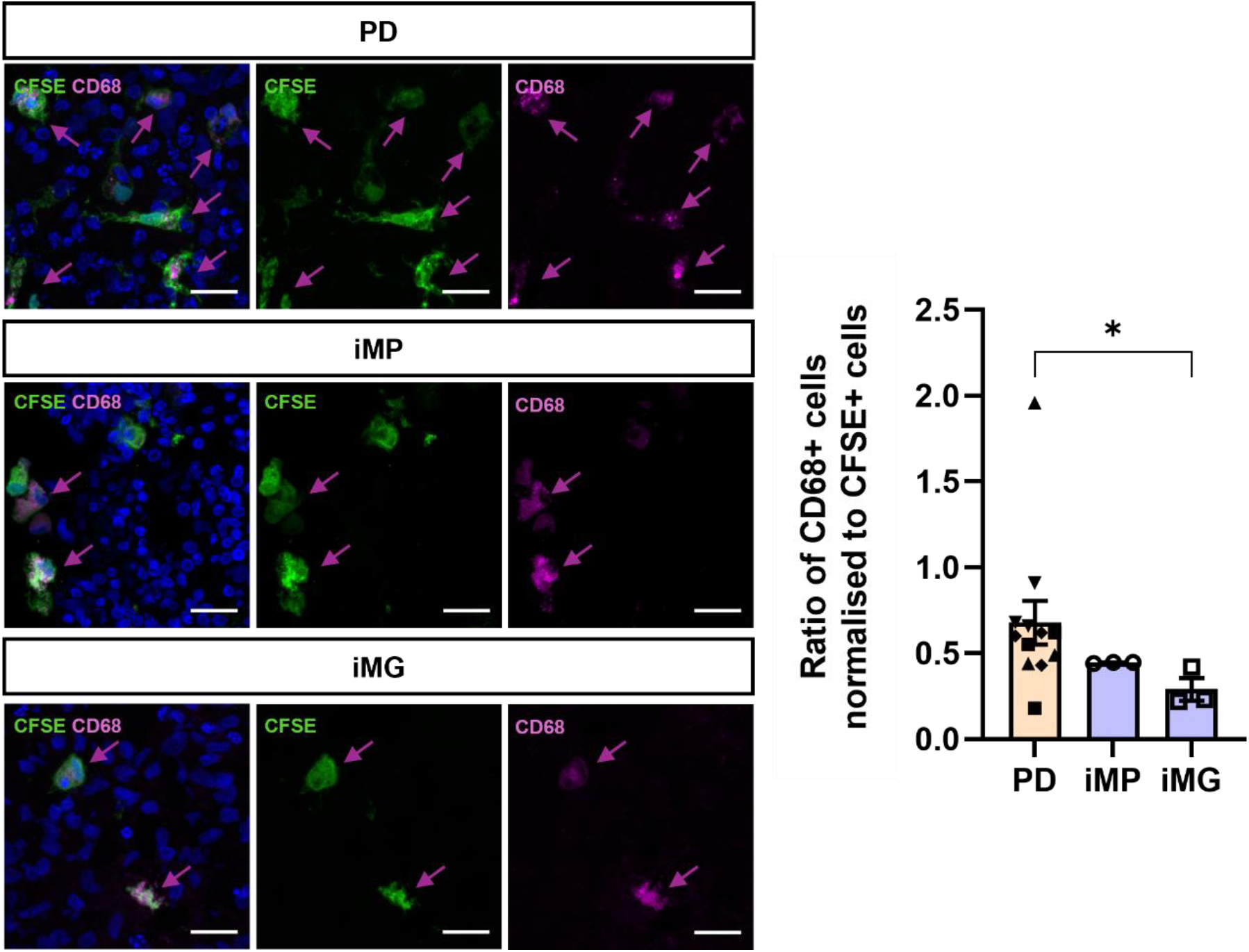
Expression of CD68 in microglia infiltrated into midbrain organoids. Representative images of infiltrated CFSE+ microglia from postmortem PD tissue (PD), iPSC-derived microglia progenitors (iMG), and iPSC-derived microglia (iMG) stained for CD68. Magenta arrows show examples of CD68+ cells. Nuclei (DAPI) in blue. Scale bars = 50 µm. Quantification of the ratio of infiltrated CFSE+ microglia that express CD68. A Kruskal-Wallis with Dunn’s multiple comparisons test was performed. Each symbol represents an organoid. Data presented as mean ±SEM. ^*^ p_adj_ ≤ 0.05 ^**^ p_adj_ ≤ 0.01 ^***^ p_adj_ ≤ 0.001.

## DISCUSSION

Microglia are important regulators for brain development, homeostasis, and neuroinflammation. These cells originate from the yolk sac during embryogenesis and subsequently migrate into the developing brain. Due to this distinct mesodermal origin, many brain organoid models naturally lack microglia and require co-culture or addition of microglia to model more accurately neuroimmune interactions. To address this issue, in this study, we utilize a 3D midbrain organoid (MO) system incorporating adult human postmortem microglia to investigate the behavior of primary human microglia in a physiologically relevant environment. This approach enables direct comparison with induced microglial progenitors (iMPs) and induced microglia (iMGs), providing insight into differences and similarities between primary and iPSC-derived microglia. To address this issue in this study, we integrate adult human postmortem microglia into midbrain organoids (MOs) that, to our knowledge, has not been previously reported. We compared the infiltration capacity and phenotypic characteristics of postmortem microglia derived from Parkinson’s disease (PD) donors with those of induced microglial progenitors (iMPs) and induced microglia (iMG). Our results show that postmortem microglia infiltrated the organoids approximately three times more efficiently and exhibited a more activated phenotype, as indicated by a higher proportion of HLA class II positive cells. In contrast, iPSC-derived microglia could have greater capacity to migrate towards the organoid core and showed higher expression of homeostatic microglial markers, including P2RY12 and IBA1.

We first examined infiltration capacity and found that all microglial groups successfully infiltrated the MOs, though their infiltration capacities varied. Postmortem PD microglia displayed approximately a three-fold higher infiltration capacity compared to the iPSC-derived cells. This enhanced infiltration of postmortem microglia might be explained by their origin from a diseased environment (Norden and Godbout, 2013). Some studies have found, after comparing microglia isolation methodologies and phenotypic states, that homeostatic signatures are often diminished after isolation due to loss of environmental cues and procedures that can induce stress and activation (Mizee et al., 2017; Rustenhoven et al., 2016). We hypothesize that this primed state may enhance the chemotactic responsiveness and ability of postmortem microglia to infiltrate the organoid tissue. In contrast, the lower infiltration of iPSC-derived cells could stem from them not being pre-conditioned to an inflammatory environment and thus possessing a lower drive to infiltrate.

Incorporation of microglia into midbrain organoids improves biological relevance by enabling neuroimmune interactions that are otherwise missing in standard organoids, supporting disease mechanism studies and recently used for toxicity testing (Sabate-Soler *et al*., 2022; Yi et al., 2025). It is important to underline the limited availability of postmortem microglia from PD patients when considering their integration into midbrain organoid models: while incorporating patient-derived microglia could greatly enhance the physiological relevance of these systems, the low tissue availability limits scalability and broad application (Mizee *et al*., 2017). As an alternative, microglia can be generated from iPSCs derived from PD patients, providing a scalable and patient-specific source of cells for such applications. Developing strategies to effectively model microglia within organoids at a high throughput scale would therefore be critical to advancing our understanding of disease mechanisms and improving the translational impact of (mid)brain organoid studies.

Following the assessment of infiltration, we investigated the phenotype of the integrated cells, starting with IBA1. Quantitative analysis revealed that while IBA1 was expressed across all groups, the proportion of IBA1^+^ cells was lower in the postmortem microglial groups compared to both iMP and iMG groups, where effectively all infiltrated cells expressed IBA1. This lower proportion of IBA1^+^ postmortem microglia could be explained by their origin from the SVZ as a previous study reported that SVZ-derived microglia express less IBA1 in mice (Ribeiro Xavier et al., 2015).

The difference in phenotype between the postmortem and iPSC-derived cells was even more pronounced when assessing activation markers. Effectively all PD microglia expressed the activation marker HLA Class II, whereas this was very low in the iPSC-derived cells. The expression of HLA Class II in postmortem microglia likely reflects the maintenance of their in vivo phenotype. The active profile of the integrated PD microglia aligns with the belief that persistent neuroinflammation, mediated by active microglia, is an important component of PD (Joers et al., 2017). The SVZ, from which the microglia were obtained, lies in close proximity to the striatum, a region profoundly affected in PD (van den Berge et al., 2013). There is a loss of dopaminergic projections from substantia nigra pars compacta that affect different regions of the brain like the striatum (Imamura *et al*., 2003). It is therefore possible that the inflammatory environment in the striatum influences the state of microglia in the adjacent SVZ, contributing to the high proportion of HLA Class II^+^ microglia that we observed.

By contrast, the low proportion of HLA Class II^+^ cells in iPSC-derived groups is consistent with their origin from a controlled in vitro environment, which is devoid of age- or disease (Haenseler et al., 2017). However, iPSC-derived microglia have been shown to express HLA class II molecules, particularly upon inflammatory stimulation, supporting their capacity to adopt antigen-presenting phenotypes in vitro (Filipello et al., 2023; Klaisner et al., 2025).

We further investigated microglial phenotype by analyzing the phagocytic marker CD68. While cells positive for this marker were observed in all groups, the proportion of CD68^+^ cells was highest in the PD group. This may suggest a phagocytic state maintained from their original disease environment, where microglia are known to clear α-synuclein aggregates and neuronal debris (Choi et al., 2020). However, this finding should be interpreted with caution since the differences in the proportion of CD68^+^ cells between the PD group and other groups were modest. Additionally, it is important to consider that these microglia are not isogenic to the organoid model, and therefore may exhibit a degree of reactivity to the host environment, which could also influence their CD68 expression.

### Limitations of the study

While this work established a novel platform for studying neuroinflammation in a 3D in vitro model by integrating postmortem microglia into MOs, it is important to discuss its methodological limitations to help guide the design of future studies that build upon this research. A primary limitation is the small sample size for several of our experimental groups. Three separate organoids were seeded per donor or iPSC-derived cell type, which for the iPSC-derived groups, could be considered a sample size of n = 3 for each condition. In addition, it should be noted that all PD donors were male. Sex differences in microglial behavior, phenotype and morphology have been observed (Han et al., 2021; Yoblinski et al., 2025). This is an important consideration, as male microglia have been reported to exhibit a more pro-inflammatory phenotype (Villa et al., 2018). The male sex of the PD donors could therefore be an additional factor contributing to the high proportion of HLA Class II^+^ cells observed in that group.

Another important limitation is that the primary microglia used in this model are not isogenic to the midbrain organoids. As a result, donor-specific differences may introduce variability, and the microglia may exhibit reactive responses to the host organoid environment. This non-isogenic nature also makes the system technically challenging to scale up or standardize, as each midbrain organoid–primary microglia pairing represents a unique donor combination. However, this variability may also be viewed as an opportunity: such a model provides a valuable platform to study primary human microglial behavior in a physiologically relevant 3D context and to compare their morphology, phenotype, and functional properties with those of iPSC-derived microglia. These comparisons could ultimately contribute to improving the culture conditions and maturation state of iPSC-derived microglia.

## CONCLUSION

This study demonstrates that integrating postmortem PD microglia could enhance the relevance of MOs for modeling neuroinflammation compared to iPSC-derived approaches. We found that postmortem microglia exhibited a more pronounced activation profile, consistent with a more mature and disease-associated state. However, the use of postmortem microglia presents important limitations. The non-isogenic nature of the model introduces additional variability, as differences between microglial donors are layered on top of variability arising from iPSC lines and organoid batch effects, requiring the inclusion of multiple donors to achieve robust conclusions. Furthermore, this approach is not readily scalable or standardizable, which limits its applicability for large-scale studies such as drug screening or systematic investigations of disease progression and mechanisms.

While logistical constraints, the model holds value as a specialized platform to study primary human microglia within a 3D neural environment, particularly for investigating their behavior, phenotype, and morphology in a disease-relevant context. As such, its application may be most appropriate for addressing focused research questions that specifically require the use of mature, primary microglia rather than for high-throughput or broadly comparative studies.

## Acknowledgments

This work was supported by a Parki Foundation prize (M.V.), a Stichting ParkinsonFonds grant (V.D.), and a Horizon ERC Advanced (ERC-AD 2022: 101097983; H.E.V.). Postmortem human brain material was obtained from the Netherlands Brain Bank (https://www.brainbank.nl/). Imaging was performed at the Microscopy and Cytometry core facility at the Amsterdam UMC.

## Author Contributions

M.V. and V.D. conceptualized project and acquired funding. M.V. and A.M.S. performed cell culture experiments. S.Z., M.V., and A.M.S. performed immunofluorescent experiments and analyses. S.Z., A.M.S., and M.V. wrote the manuscript. A.D. provided feedback on the study design and on the manuscript. H.E.V. and V.D. provided resources and feedback on the manuscript. All authors revised and approved the final manuscript.

## Supplemental MATERIALS AND METHODS

### Human postmortem brain tissue collection & microglia isolation

Fresh postmortem dorsal subventricular zone (SVZ) tissue was obtained from four Parkinson’s disease (PD) donors (n = 4) through the Netherlands Brain Bank (NBB). The donors were males aged 74, 67, 75, and 82 years, with one donor having combined Dementia with Lewy Bodies (DLB) and PD. Microglia were isolated from the SVZ tissue using magnetic cell separation (MACS; Miltenyi Biotec) with anti-CD11b-coated microbeads, as described in(Donega et al., 2019). Following isolation, CD11b^+^ microglia were cryopreserved in StemCellBanker (Amsbio, 11924).

### iPSC culture

In this study, the healthy control iPSC line TMOi001 (Gibco, A18945) was used. The iPSCs were cultured feeder-free on Matrigel-coated (Corning, 356234) culture dishes in mTeSR plus medium (STEMCELL Technologies, 100-0276) at 37 °C with 5% CO2. The iPSCs were passaged once a week (1:50) with 0.5 mM ethylenediaminetetraacetic acid (EDTA; Promega, 6381-92-6). Following passaging, the medium was supplemented with 5 µM ROCK-inhibitor Y-27632 (RI; SigmaAldrich, SCM075) for 24 hours (hrs).

### Generation of Midbrain Organoids (MOs)

NPCs were generated according to Reinhardt et al., 2013, with minor adaptations. Briefly, single cell iPSCs were seeded in ultra-low attachment round-bottom 96-well plates in human embryonic stem cell medium (DMEM/F-12 [Gibco, 31330-038], 20% Knock-Out Serum Replacement [ThermoFisher, 10828028], 1% GlutaMAX [Gibco, 2360312], 1% Non-Essential Amino Acids [Gibco, 2465992], 1% Penicillin-Streptomycin [P/S; Gibco, 15070063], and 0.0007% β-mercaptoethanol [Gibco, 31350]) with patterning factors (Reinhardt et al., 2013). By day 2, formed embryoid bodies were transitioned to N2B27 medium, composed of a 1:1 mixture of DMEM/F-12 + GlutaMAX (Gibco, 31331028) and Neurobasal (ThermoFisher, 21103049), with 1% P/S, 1% B27 without Vitamin A (Gibco, 12587010) and 0.5% N2 (ThermoFisher, 17502001). Differentiation was then guided by time-dependent combinations of small molecules to drive commitment to the NPC fate (Reinhardt *et al*., 2013). NPCs were maintained and passaged weekly. On day 37, NPCs were cryopreserved in N2B27 medium with small molecules and 10% Dimethyl Sulfoxide (Sigma-Aldrich, D2650).

MOs were generated according to Monzel et al. 2017. Briefly, single cell NPCs were seeded in an ultra-low attachment round bottom 96-well plate at a density of 18,000 cells per well. From DIV2 to DIV31, the medium was supplemented with patterning and maturation factors (Monzel *et al*., 2017). From DIV31, no factors were added to the medium.

### Generation of human iPSC-derived microglial progenitors and mature microglia

Microglial cells were differentiated from hiPSCs as described in (Kenkhuis et al., 2022; Sabogal-Guáqueta et al., 2023) with minor adaptations. hiPSCs were detached using accutase (Stemcell Technologies, 07920) and re-suspended in mTeSR PLUS medium supplemented with 10 µM Y-27632, 10 µg/ml recombinant human BMP4 (Thermo Fisher, PHC9534), 10 µg/ml recombinant human VEGF (PeproTech, 100-20), and 10 µg/ml recombinant human SCF (ThermoFisher, PHC2115). 4 x 10^6^ cells from the single-cell suspension were plated per well of AggreWelltm 800 (StemCell Technologies, 34815), and the plate was centrifuged to form mesodermal embryoid bodies (mEB). Three-quarters of the media were refreshed daily for 6 days. The mEBs were plated in X-VIVO 15 medium (Lonza, BE02-060Q) supplemented with 100 U/mL penicillin-streptomycin (Invitrogen, 1514012), 2 mM Glutamax (ThermoFisher, 35050-038), 50 µM 2-β-mercaptoethanol (ThermoFisher, 31350-01), 100 µg/ml recombinant human M-CSF (Peprotech, 300-25), and 25 µg/ml recombinant human IL-3 (Peprotech, 200-03). Half of the media was refreshed weekly for 3-4 weeks and once mEBs started producing myeloid precursor cells they were harvested weekly. Microglial progenitors were harvested from the supernatant, resulting in iMPs. The remaining harvested floating cell population was collected, passed through a 40μm cell strainer to ensure single-cell suspension, and pelleted via centrifugation (300g for 5 minutes). Cells to become iMGs were plated for two weeks in Advanced DMEM/F12 (ThermoFisher, 12634-010) supplemented with 2 mM Glutamax, 100 U/mL Pen-Strep, 55 uM 2-mercaptoethanol and 100 ng/m IL-34 (Peprotech, 200-34), 25 ng/mL M-CSF (Peprotech, 300-25), 50 ng/mL TGF-B1 (Peprotech, 100-21C) and 10ng/ml GM-CSF (PeproTech, 300-03). During this period, the media was refreshed every 2 days by performing a half-medium exchange

### Microglial integration into MOs

#### Preparation of human postmortem microglial for co-culture

Postmortem microglia were thawed under warm tap water and collected in DMEM/F-12 + GlutaMAX and 10% Fetal Bovine Serum (ThermoFischer, A31605). Cells were labelled using the CellTrace Carboxyfluorescein succinimidyl ester (CFSE) Cell Proliferation Kit (ThermoFischer, C34570) at a concentration of 1 μM and incubated for 20 min at 37 °C. Reaction was quenched with DMEM and 10% Fetal Bovine Serum (FBS). CFSE-labelled microglia were resuspended in N2B27 medium immediately prior to seeding.

#### Preparation of iMP and iMG for co-culture

iMP and iMG were labelled with CFSE as described above. CFSE-labelled iMP were resuspended in a 1:1 mixture of N2B27 and Microglia Maturation medium. Microglia Maturation medium was composed of Advanced DMEM/F-12 (ThermoFisher, 12634-010) with 1% P/S (Invitrogen, 15140122), 1% GlutaMAX (ThermoFisher, 35050-038), 0.1% 2-mercaptoethanol (ThermoFisher, 31350-010), 1% N2 supplement (ThermoFisher, 17502-048), and the following recombinant human cytokines: 10 ng/mL GM-CSF (PeproTech, 300-03), 100 ng/mL IL-34 (Peprotech, 200-34), 50 ng/mL TGF-β1 (Peprotech, 100-21C), and 25 ng/mL M-CSF (Peprotech, 300-25). CFSE-labelled iMG were resuspended in N2B27 medium immediately prior to seeding.

#### Seeding and co-culture maintenance

To create co-cultures, MOs were seeded on DIV36 with either human postmortem microglia from one of the PD donors, iMP, or iMG. Each MO was seeded with 30,000 microglia and maintained in co-culture for 14 days. For each condition, 3 MOs were seeded with the same microglial suspension as biological replicate, resulting in 12 MOs in the postmortem group, 3 MOs in the iMP group, and 3 MOs in the iMG group. Co-cultures with postmortem microglia or iMG were maintained in N2B27 medium. Those with iMP were maintained in a 1:1 mixture of N2B27 and Microglia Maturation medium. Medium was refreshed every 2-3 days. On day 14 of co-culturing (DIV50), MOs were fixed with 4% paraformaldehyde for 1 hr at 4 °C.

A fraction of microglia were plated as mono-culture on an 8-well chamber slide (Ibidi, 80806) pre-coated with 0.01 mg/mL Poly-D-Lysine (Sigma-Aldrich, P0899-50MG) and 0.01% gelatin (Sigma-Aldrich, G1393-20M) and fixed the next day with 4% PFA for 20 min 4 °C.

### Immunofluorescence staining and analysis

#### 2D staining

After overnight incubation in 30% sucrose at 4 °C, MOs were snapfrozen in OCT mounting medium (VWR, 361603E) and cut on a cryostat (ThermoFischer) into 20 µm thick sections, which were mounted on Superfrost Plus adhesion slides (Epredia, J1800AMNZ). Slides were thawed and washed in PBS with 0.05% Tween-20 (PBS-Tween; Sigma-Aldrich, P7949) for 3 × 10 min. Blocking was performed using 3% Bovine Serum Albumin (Roche, 10 735 086 001) and 1% Triton-X100 (Merck, 8603) in PBS for 1 h at RT. Antibodies (**Table S1**) were diluted in blocking solution. Primary antibody was incubated overnight at 4 °C and secondary antibodies with DAPI (1:2000, Sigma-Aldrich, D9542) 1 hr at RT. Sections were mounted in MOWIOL 4-88 (Sigma-Aldrich, 81381-250G) containing 2.5% DABCO (Sigma-Aldrich, 290734).

High resolution images were obtained using a Nikon Ti2 AX confocal microscope equipped with a 40x APO LWD λS 40× water immersion objective (NA = 1.15, WD = 610 µm). Imaging was performed using Galvano scanning with 2x line averaging (0.084 fps), a pixel size of 0.109 µm and a pinhole size of 1.2 AU. Maximum intensity projections were generated from z-stacks with a z-step size of 1 µm, with total stack depths ranging from approximately 8 µm to 13 µm across images. Entire organoid sections were imaged at 40x magnification on an Olympus VS200 slide scanner equipped with a 0.95 NA air/dry UPLXAPO objective and a VS304M camera, at a resolution of 0.1725 µm/pixel. With VS200 ASW software (version 3.3), maximum intensity projections were generated from 14 µm z-stacks with a z-step size of 2 μm. Multiple sections per organoid were imaged an subsequently pooled per organoid.

Image analysis was performed in QuPath (version 0.6.0). Nuclei were detected per image using the StarDist2D extension, with a size between 10 µm^2^ and 200 µm^2^ and minimum mean DAPI intensity of 100. Machine learning–based object classification was conducted for each marker and the percentage of positive cells per total DAPI+ cells was calculated. To account for differences in number of microglial cells, the percentage of positive cells was normalized to the average percentage of CFSE+ cells per total DAPI+ cells per experimental group.

#### 3D staining

Whole organoids were stained with the following steps: (1) permeabilization with 2% Triton-X100 in PBS for 24 hrs at 4 °C, (2) blocked with 3% Normal Donkey Serum and 0.3% Triton-X100 in PBS, (3) antibody incubation in blocking solution for 48 hrs at 4 °C on an orbital shaker for both primary and secondary antibodies, (4) incubation with RapiClear 1.49 Solution (#RC149001) for 72 hrs at RT. Whole MOs were scanned with a Nikon Ti2 AX confocal microscope equipped with a PLAN APO λD 10x OFN25 DIC N1 (NA = 0.45, WD = 4000µm) or PLAN APO λD 20x OFN25 DIC N2 (NA = 0.80,

WD = 800µm). With 10x objective, imaging was performed using Resonant scanning (0.48 fps), pixel size of 0.539 µm, and pinhole size of 1.2 AU. Maximum intensity projections were generated from z-stacks with a z-step size of 2 µm. With the 20x objective, imaging was performed using Galvano scanning (0.229 fps), pixel size of 0.157 µm, and pinhole size of 1.2 AU. Maximum intensity projections were generated from z-stacks with a z-step size of 1 µm.

**Table S1.** Antibodies used in this study.

| Antibody | Host species | Final dilution | Manufacturer | Catalog number |
| --- | --- | --- | --- | --- |
| CD68 | Mouse | 1:500 | DAKO | M0814 |
| Fluorescein | Rabbit | 1:1000 | Abcam | ab19491 |
| HLA Class II | Mouse | 1:500 | DAKO | M0775 |
| IBA1 | Rabbit | 1:500 | WAKO Chemicals USA | 019-19741 |
| P2RY12 | Rabbit | 1:1000 | Invitrogen | 702516 |
| Donkey anti-mouse Alexa Fluor 555 | Donkey | 1:2000 | Invitrogen | A31570 |
| Donkey anti-mouse Alexa Fluor Plus 647 | Donkey | 1:1000 | Invitrogen | A32787 |
| Donkey anti-rabbit Alexa Fluor 488 | Donkey | 1:2000 | Invitrogen | A21206 |
| Donkey anti-rabbit Alexa Fluor Plus 647 | Donkey | 1:1000 | Invitrogen | A32795 |

### Statistical analysis

Data visualization and statistical analyses were performed using GraphPad Prism (version 10.2.0). Per experimental group, outliers in organoid sections were determined with the ROUT method (Q = 1%) before calculating the mean per organoid. Normal distribution was assessed with Shapiro-Wilk test. Normally distributed data were analyzed using one-way ANOVA with Tukey’s multiple comparisons test (P2RY12, IBA1, HLA). Non-normally distributed data were analyzed using a Kruskal-Wallis with Dunn’s multiple comparisons test (CFSE, CD68). An adjusted p-value ≤ 0.05 was considered statistically significant. All data are depicted as mean ± SEM.

